# Kinetic mechanisms of fast glutamate sensing by fluorescent protein probes

**DOI:** 10.1101/664458

**Authors:** C. Coates, S. Kerruth, N. Helassa, K. Török

**Affiliations:** St. George’s, University of London

**Keywords:** glutamate, biosensor, fluorescence

## Abstract

Protein-based fluorescent glutamate sensors have the potential for real-time monitoring of synaptic and cellular glutamate concentration changes, however even the fastest currently available sensors’ response times of 2-3 ms are too slow for accurate reporting of the post-synaptic AMPA receptor function in physiological conditions. We have developed probes based on the bacterial periplasmic glutamate/aspartate binding protein with either an endogenously fluorescent protein or a synthetic fluorophore as the indicator of glutamate binding: affinity variants of iGluSnFR termed iGlu_*h*_, iGlu_*m*_ and iGlu_*l*_ covering a range of *K*_d_−s (5.8 μM, 2.1 mM and 50 mM, respectively) and a novel fluorescently labelled indicator, Fl-GluBP with a *K*_d_ of 9.7 μM are presented. The fluorescence response kinetics of all the probes are consistent with two-step mechanisms involving ligand binding and rate limiting isomerisation, however the contribution in each step to the total fluorescence enhancement and kinetic paths to the final state are diverse. In contrast to the previously characterised ultrafast indicators iGlu_*u*_ and iGlu_*f*_, for which fluorescence enhancement occurred only in the rate limiting isomerisation step, the sensors described here all have biphasic binding kinetics with a significant fraction of the fluorescence increase evoked by glutamate binding which, in the case of iGlu_*h*_ and Fl-GluBP, occurs with a diffusion limited rate constant. The above genetically encoded and chemically labelled fluorescent glutamate sensor variants demonstrate how single amino acid changes around the binding site introduce structural heterogeneity affecting the kinetic mechanism of interactions with glutamate. Through their broad affinity range and mechanistic variety, the probes contribute to a novel toolkit for monitoring processes of glutamate neurotransmission and cellular homeostasis.

**STATEMENT OF SIGNIFICANCE:** Glutamate is a major excitatory neurotransmitter, important in synaptic plasticity e.g. memory formation. Although predicted to clear rapidly from the synaptic cleft following presynaptic release, optical monitoring of glutamate neurotransmission has only become possible with the advent of fluorescent, protein-based indicators. Understanding their biophysical properties is important for quantification of the observed processes. Here we report the biophysical characterisation of a number of glutamate indicator variants based on the bacterial periplasmic glutamate/aspartate binding protein, revealing the subtle differences in their kinetic pathway caused by structural alteration of the glutamate binding protein by point mutations. Diffusion limited glutamate binding indicated by a novel chemically labelled probe hints at the mechanism that underlies the rapid response of the AMPA receptor.

## INTRODUCTION

Glutamate is a major excitatory neurotransmitter in the central nervous system, however its synaptic and cellular dynamics have only become possible to investigate with high spatial and temporal resolution with the emergence of well-functioning optical sensors (1,2). The fastest fluorescent glutamate sensors iGluSnFR variants iGlu_*f*_ and iGlu_*u*_ have proved useful for tracking high frequency glutamate release at single hippocampal synapses (3) and revealed impaired glutamate clearance at striatal synapses in HD mice (4).

Information processing at synapses is rapid: glutamate neurotransmitter release is fast evoking AMPA receptor (AMPAR) channel opening with a time constant, *τ*_on_ of 17 μs (5), making AMPAR the fastest responding ligand-gated ion channel. Glutamate clearance from the synapse is predicted to occur with *τ*_off_ of 50-200 μs (6,7). Visualising synaptic and intracellular glutamate offers an important approach for better understanding of mechanisms of information processing at the synapse and of cellular glutamate homeostasis. Real-time tracking of neurotransmitter release requires rapid response sensors and for the investigation of glutamate homeostasis, sensors with affinities corresponding to physiological conditions are required.

Fluorescent glutamate sensors were initially developed from the extracellular domains of AMPAR, constructs of which are termed S1S2 and later from the bacterial periplasmic glutamate/aspartate binding protein (GluBP). GluR2-AMPAR-derived S1S2 constructs labelled with synthetic fluorophores were promising candidates for the generation of a glutamate biosensor, but have proved impractical due to low refolding yield and stability (8). Following extensive engineering, stable fluorescent S1S2 derivatives with high fluorescence dynamic ranges have been reported (9,10). For a fluorescent S1S2 derivative eEOS, an association rate constant of 1.2 × 10^5^ M^−1^s^−1^ and an *off*-rate of 14 s^−1^ were measured at 25 °C (10), too slow for tracking synaptic glutamate dynamics.

Bacterial periplasmic ligand binding proteins have been widely used for the generation of fluorescent biosensors, initially by covalent derivatisation with synthetic fluorophores, e.g. for inorganic phosphate (11) and amino acid and sugar ligands (12,13). Ligand binding kinetics of members of the bacterial periplasmic ligand binding protein family have been measured by monitoring Trp fluorescence, revealing association rate constants in the order of 10^7^ M^−1^s^−1^ (14). The AMPAR S1S2 domain shares structural homology with GluBP resulting in similar glutamate binding kinetic parameters measured by Trp fluorescence changes (15). The mechanism derived from these investigations was termed the Venus flytrap whereby slow ligand binding is followed by a rapid conformational change, representing domain closure to trap the bound ligand. However, as Trp fluorescence served as an indicator of the conformational change, not of the binding, these experiments did not reveal the true association rate constant for glutamate.

In the genetically encoded glutamate sensor iGluSnFR, two separated fragments of GluBP were fused with circularly permuted (cp) EGFP. Fluorescence enhancement is based on two flanking portions of GluBP (large fragment GluBP 1-253, depicted as iGlu_l_ and small fragment GluBP 254-279, depicted as iGlu_s_) non-covalently reattaching on glutamate binding and thereby correcting the structure of cpEGFP (**Fig. 1A**). Apo-iGluSnFR has low fluorescence; to achieve a highly fluorescent state, reconstitution of GluBP is required, stabilised by bound glutamate. The kinetic mechanism we have previously determined, is illustrated on the example of the ultrafast sensor iGlu_*u*_. Its GluBP 1-253 fragment first binds glutamate, this is however not sufficient for fluorescence enhancement. Binding is followed by a conformational change (the reattachment of GluBP 254-279 fragment to form the complete structure, depicted as iGlu_c_) during which the highly fluorescent state develops. The rate of this isomerisation step limits the fluorescence response (**Scheme 1**).

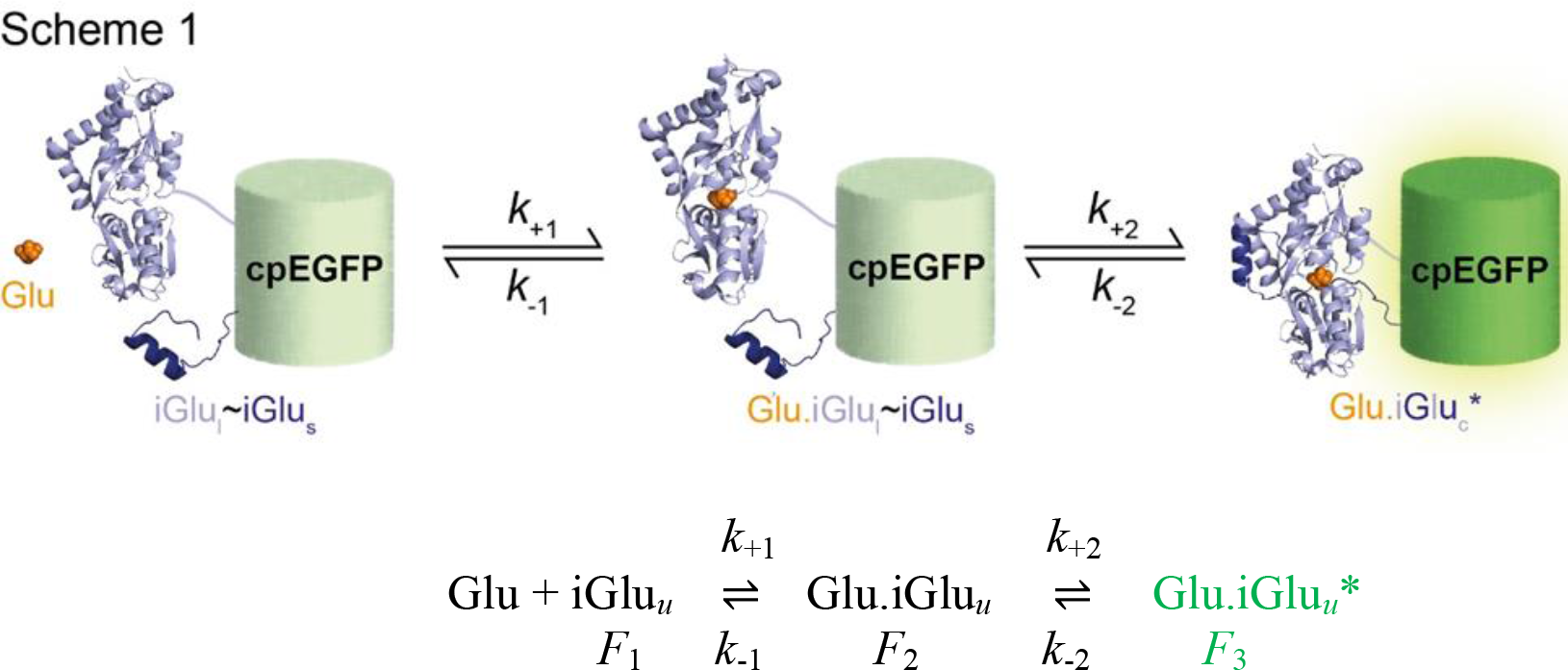

**Figure 1.**
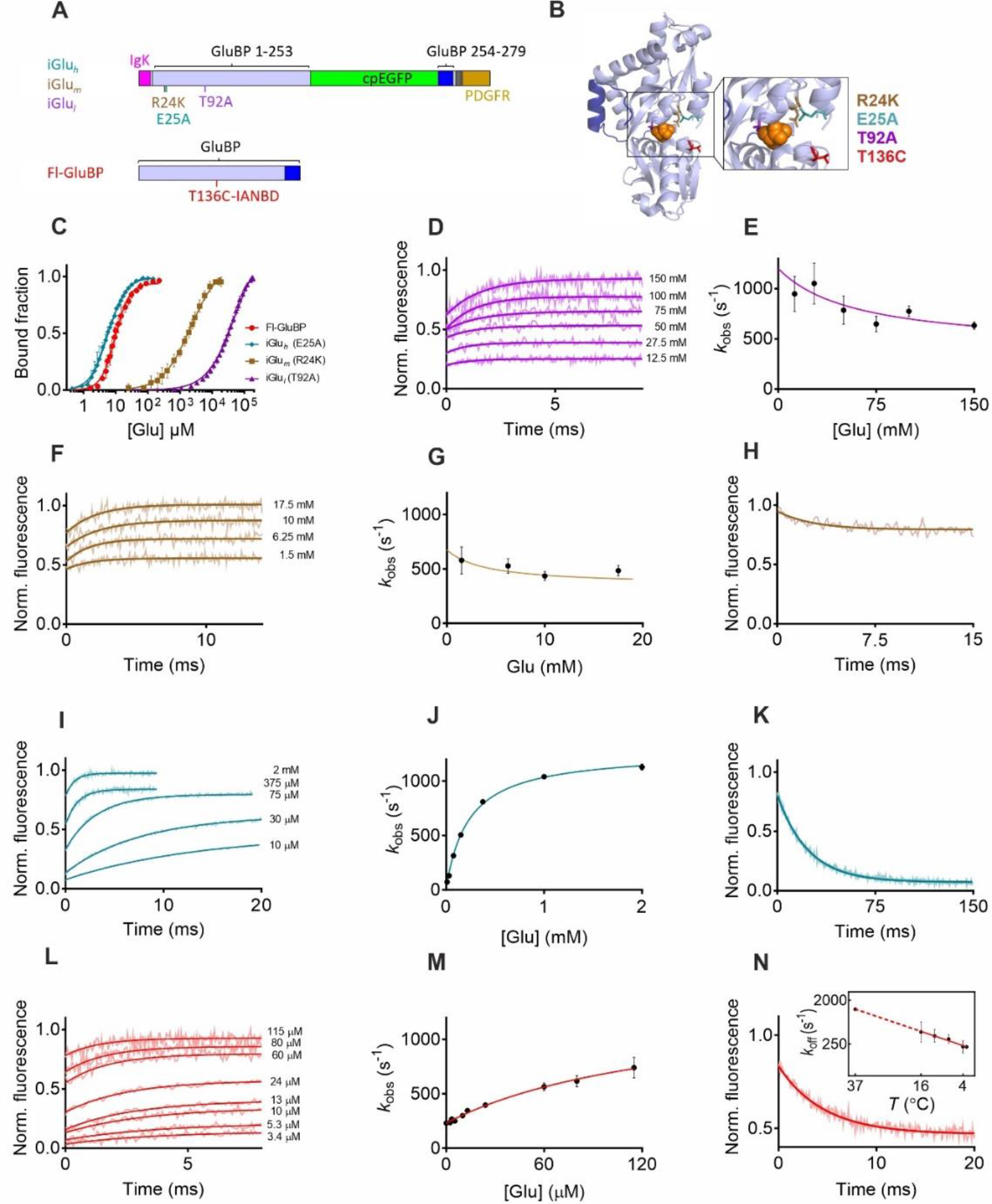
Design and biophysical properties of iGlu variants and Fl-GluBP. (**A**) Domain structure and labelling site. (**B**) Crystal structure of GluBP (PDB ID: 2VHA) with mutation sites indicated. (**C**) Equilibrium titration with glutamate of iGlu variants and Fl-GluBP (20 °C). (**D**) Association kinetic records of iGlu-T92A (iGlu_*l*_) (20 °C). (**E**) Plot of observed association rates (*k*_obs_) against glutamate concentration for iGlu_*l*_. (**F**) Association kinetic records of iGlu-R24K (iGlu_*m*_) (20 °C). (**G**) Plot of observed association rates (*k*_obs_) against glutamate concentration for iGlu_*m*_. (**H**) Dissociation kinetics of iGlu_*m*_ (20 °C). (**I**) Association kinetic records of iGlu-E25A (iGlu_*h*_) (20 °C). (**J**) Plot of observed association rates (*k*_obs_) against glutamate concentration for iGlu_*h*_. (**K**) Dissociation kinetics of iGlu_*h*_ (20 °C). (**L**) Association kinetic records of Fl-GluBP (3 °C). (**M**) Plot of observed association rates (*k*_obs_) against glutamate concentration for Fl-GluBP. (**N**) Dissociation kinetics of Fl-GluBP (3 °C). Inset: Arrhenius plot of temperature dependence of the *off*-rate.

The single exponential fluorescence enhancement observed for iGluSnFR and fast variants iGlu_*f*_ and iGlu_*u*_ thus corresponds to the glutamate binding induced conformational change which reflects the two spatially separated fragments of GluBP recombining allowing the cpEGFP β-barrel to seal. To-date, the fastest sensor iGlu_*u*_ has *τ*_on_ of 460 μs in solution and *τ*_off_ of 2.7 ms (34 °C) at Shaffer collaterals in hippocampal slices(3).

We hypothesized that diffusion limited glutamate binding occurs but is not indicated by the already characterised probes and hence here we explore the kinetic mechanism of a selection of affinity variants to see if they reveal rapid glutamate binding kinetics. We find that the kinetic mechanisms of novel iGluSnFR affinity variants (low affinity iGlu_*l*_, medium affinity iGlu_*m*_ and high affinity iGlu_*h*_) are diverse, suggesting structural heterogeneity introduced by the binding site mutations. Furthermore, a novel sensor Fl-GluBP, generated by targeted Cys substitution of GluBP and derivatisation with a synthetic fluorophore, displays diffusion limited glutamate association kinetics. Such a sensor has the potential for real time glutamate tracking at single synapses under high frequency stimulation.

## MATERIALS AND METHODS

### Materials

pET41a iGlu_*m*_ (R24K), pET41a iGlu_*h*_ (E25A), pET41a iGlu_*l*_ (T92A) and pET30b GluBP were generated as previously described (3) and are available on ADDGENE (Watertown, MA) (119829, 119830, 119832 and 119835, respectively). *Escherichia coli* XL10-Gold and BL21(DE3) Gold cells were purchased from STRATAGENE (San Diego, CA). 6-Acryloyl-2-Dimethylaminonaphthalene (Acrylodan), 7-Diethylamino-3-(4’-maleimidylphenyl)-4-methylcoumarin (CPM), N-((2-(iodoacetoxy)ethyl)-N-Methyl)- amino-7-Nitrobenz-2-Oxa-1,3-Diazole (IANBD ester) and Oregon Green 488 maleimide were purchased from LIFE TECHNOLOGIES LTD (Paisley, UK) and 6-bromoacetyl-2-dimethylaminonaphthalene (BADAN) from EUROGENTEC (Southampton, UK). N-(2-(iodoacetamido)ethyl)-7-diethylaminocoumarin-3-carboxamide (IDCC) was a gift from J.E.T. Corrie, NIMR, London.

### Site-directed mutagenesis

Ser or Thr to Cys mutations were introduced into pET30b GluBP via site-directed mutagenesis according to the QuikChange II XL protocol (AGILENT TECHNOLOGIES, Santa Clara, CA) using the following primers:

T71C 5’-GCAGGTAAAACTGATTCCGATTTGCTCACAAAACCGTATTCC-3’
S72C 5’- TAAAACTGATTCCGATTACCTGCCAAAACCGTATTCCACTGCTG-3’
T83C 5’-CCACTGCTGCAAAACGGCTGTTTCGATTTTGAATGTGGTTC-3’
S90C 5’- ACTTTCGATTTTGAATGTGGTTGTACCACCAACAACGTC-3’
T91C 5’- CGATTTTGAATGTGGTTCTTGCACCAACAACGTCGAACGC-3’
T92C 5’- GATTTTGAATGTGGTTCTACCTGCAACAACGTCGAACGCC-3’
T136C 5’-CAAAGCCGTAGTCGTCTGTTCCGGCACTACCTCTG-3’
S137C 5’-CGTAGTCGTCACTTGCGGCACTACCTCTGAAG-3’
T140C 5’-GTCGTCACTTCCGGCACTTGCTCTGAAGTTTTGCTCAAC-3’
A210C 5’- GCCGCAGTCTCAGGAGTGCTACGGTTGTATGTTG-3’

DNA sequences were verified by sequencing (GENEWIZ UK LTD, Bishop’s Stortford, UK).

### Protein expression and purification

iGluSnFR variants (iGlu_*l*_, iGlu_*m*_ and iGlu_*h*_) and GluBP proteins (GluBP 600n and GluBP-T136C) were expressed and purified as previously described (3). Briefly, cells were grown at 37 °C until OD_600nm_ 0.5-1.0 and expression was induced with 0.4 mM IPTG, overnight at 20 °C. Cells were recovered by centrifugation and lysed by sonication on ice. Clarified lysates were purified by affinity purification (GSTrap or HisTrap, GE HEALTHCARE, Little Chalfont, UK) and purity was assessed by SDS-PAGE. Purified proteins were dialysed overnight at 4 °C in 50 mM HEPES-Na^+^, 200 mM NaCl pH 7.5 and stored at −80 °C.

### Protein labelling with thiol-reactive environmentally-sensitive fluorophores

Purified GluBP-T136C was labelled overnight at 4 °C using a 2-fold excess of fluorophore (Acrylodan, IDCC, CPM, Oregon Green 488 Maleimide, BADAN or IANBD). Labelled protein was then dialysed three times over a 24-hour period at 4 °C against 500 volumes of PBS to remove unreacted dye, then once against 300 volumes of assay buffer for buffer exchange.

### Measurement of protein concentration

Protein concentration was determined by UV spectroscopy. For GluBP T136C-IANBD (Fl-GluBP), an extinction coefficient of 25,000 M^−1^ cm^−1^ at 495 nm was used, which corresponds to the absorbance peak of IANBD ester. For His-GluBP proteins and iGluSnFR variants (iGlu_*h*_, iGlu_*m*_ and iGlu_*l*_) the extinction coefficients were and 24,075 M^−1^ cm^−1^ and 90,690 M^−1^ cm^−1^ respectively, at 280 nm (16).

### Dynamic range and affinity measurements

Measurements were carried out using a Fluorolog3 spectrofluorimeter (HORIBA UK LTD, Northampton, UK). For dynamic range determination, fluorescence emission spectra were recorded in the presence and absence of ligand. *F*_+ligand_/*F*_−ligand_ was calculated using values at the fluorescence emission maximum. For determination of ligand affinity and specificity (glutamate, aspartate, glutamine), an appropriate stock solution of ligand was added to the glutamate sensor (iGlu variant or labelled-GluBP) either manually or via continuous titration (10 μL/min) using an automated syringe pump (ALADDIN 1000, WPI, Hitchin, UK). Measurements were performed in a stirred 3 mL cuvette containing protein in assay buffer (50 mM HEPES-NaOH pH 7.5, 100 mM NaCl, 2 mM MgCl_2_). For iGluSnFR variants λ_ex_ = 492 nm and λ_em_ = 512 nm, for Fl-GluBP λ_ex_ = 495 nm and λ_em_ = 535 nm. Data were corrected for dilution, normalised and the dissociation constant (*K*_d_) and Hill coefficient (*n*, when appropriate) were determined by fitting the data using the ‘one site specific binding’ equation (with or without Hill slope as appropriate) in GraphPad Prism 7. Titrations were performed at least in triplicates and the error expressed as mean ± SD.

### Stopped-flow fluorimetry

Experiments were performed as detailed in (3) Helassa *et al.* (2018). Briefly, a KinetAsyst SF-61DX2 system (TGK SCIENTIFIC, Bradford on Avon, UK) equipped with a temperature manifold (17) was used to measure kinetics. Fluorescence excitation was set to 492 nm and fluorescence emission was collected using a long pass filter (>530 nm). For association kinetics, 0.25 – 0.7 μM protein (concentrations in the mixing chamber) was rapidly mixed with increasing concentrations of glutamate. For dissociation, GluBP 600n was used in excess (915 – 1200 μM) and rapidly mixed with glutamate-bound sensor. η values were calculated for 3 °C using the density and viscosity calculator for glycerol/water mixtures based on Cheng (2008) at: http://www.met.reading.ac.uk/~sws04cdw/viscosity_calc.html.

Data shown is the average of at least four traces, data was then fit using single or double exponentials (as appropriate) to ascertain the fluorescence rise or decay rate, using KinetAsyst software (TGK SCIENTIFIC, Bradford on Avon, UK).

## RESULTS

Here we investigate the kinetic mechanisms of genetically encoded affinity variants termed iGlu-T92A (iGlu_*l*_), iGlu-R24K (iGlu_*m*_), iGlu-E25A (iGlu_*h*_) and chemically labelled Fl-GluBP (Fig. 1A,B).

First the equilibrium glutamate binding and fluorescence properties are presented. iGlu-T92A (iGlu_*l*_) has the lowest affinity (*K*_d_ of 50 ± 2 mM) with glutamate induced fluorescence enhancement *F*_(+Glu)_/*F*_(−Glu)_ of 1.7 ± 0.5. iGlu-R24K (iGlu_*m*_) is the medium affinity probe (*K*_d_ of 2.1 ± 0.1 mM) with *F*_(+Glu)_/*F*_(−Glu)_ of 2.5 ± 0.4. iGlu-E25A (iGlu_*h*_) has the highest affinity for glutamate (*K*_d_ of 5.8 ± 0.2 μM) with *F*_(+Glu)_/*F*_(−Glu)_ of 3.4 ± 0.6, of the genetically encoded variants. We furthermore present Fl-GluBP (*K*_d_ of 9.7 ± 0.3 μM, *F*_(+Glu)_/*F*_(−Glu)_ of 2.9 ± 0.1) generated by labelling GluBP with environmentally sensitive fluorophore IANBD near the binding site, substituting selected amino acids for Cys (**Fig. 1C** and **Table 1**). The kinetic mechanisms of each of the four glutamate sensors will be presented in turn revealing three different kinetic pathways.

**Table 1.**
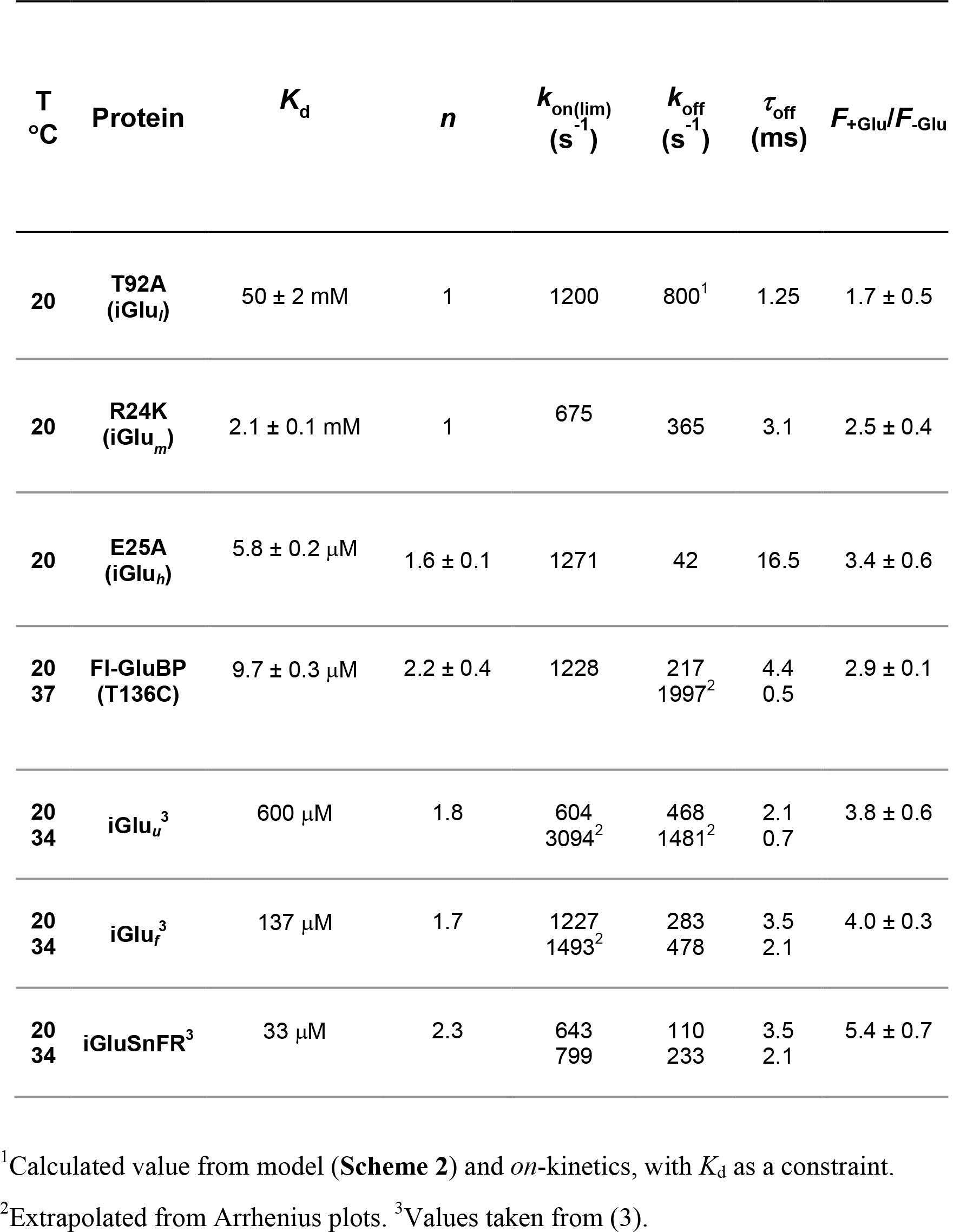
Kinetic parameters of iGlu_*l*_, iGlu_*m*_, iGlu_*h*_ and Fl-GluBP obtained by measurement

### Kinetic mechanism of low and medium affinity sensors iGlu-T92A (iGlu_l_) and iGlu-R24K (iGlu_m_)

Both iGlu-T92A (iGlu_*l*_) and iGlu-R24K (iGlu_*m*_) showed biphasic kinetics: a fast fluorescence increase with rates too fast to measure was followed by a single exponential fluorescence rise on glutamate binding with observed rates of up to 1200 s^−1^ (**Fig. 1D,E**) and 675 s^−1^ (**Fig. 1G,H**). For both probes the observed association rate, *k*_obs2_ decreased as [glutamate] was increased, plateauing at 400 s^−1^ and 350 s^−1^, respectively. Such pattern of the association rate plot is consistent with a mechanism in which a slow pre-equilibrium exists between two forms of the apo-protein, when glutamate binds to one of them preferentially (**Scheme 2**). On the example of iGlu_*l*_, iGlu_l_-iGlu_s_(T92A) and iGlu_c_(T92A) are in equilibrium, glutamate binds to iGlu_c_(T92A) preferentially. iGlu_c_(T92A) has greater fluorescence intensity than iGlu_l_-iGlu_s_(T92A) and glutamate binding results in a further fluorescence enhancement. The same description applies to iGlu-R24K (iGlu_*m*_) as shown in **Scheme 2**. Dissociation kinetics were measured by trapping released glutamate from the complex of iGlu_*l*_ or Glu_*m*_ with glutamate with > 100-fold excess of purified GluBP 600n (see **Materials and Methods**). While dissociation kinetics were not possible to measure for iGlu_*l*_ given the combination of its low affinity and fluorescence dynamic range, a single exponential fluorescence decay was obtained for Glu_*m*_ with a rate of 365 ± 58 s^−1^ (20 °C) (**Fig. 1H**). The lack of an initial fast phase and the only a partial fluorescence decrease in the dissociation record indicated that significant rebinding of glutamate occurred in the conditions used. Data best fitted to **Eq. 1** (18), giving the following set of parameters: for iGlu_*l*_, *K*_d4_, 60 mM; *k*_+3_, 400 s^−1^ and *k*_−3_, 800 s^−1^. The calculated *K*_d(overall)_ of 40 mM is in good agreement with the measured *K*_d_ of 50 ± 2 mM. Final relative fluorescence *F*_∞_ was 1.7. Relative fluorescence values for *F*_1_, *F*_2_ and *F*_3_ are 0.86, 1.28 and 1.7, respectively were calculated from **eq. 2** (19). For Glu_*m*_, best fit values from the rate plot are *K*_d4_, 4 mM; *k*_+3_, 350 s^−1^ and *k*_−3_, 325 s^−1^. The *K*_d(overall)_ calculated from the fitted values is 1.9 mM, which is in good agreement with the measured value of 2.1 ± 0.1 mM. Gutamate binding to iGlu_c_(R24K) gives *F*_∞_ = 2.5. Using **eq. 2**, *F*_1_ of 0.68 is calculated for iGlu_l_-iGlu_s_(R24K); for the apo-form iGlu_c_(R24K) that glutamate preferentially binds to, *F*_2_ of 1.345 is obtained and *F*_3_ for iGlu_c_(R24K)* is 2.5 (**Table 2**).

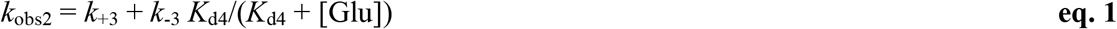

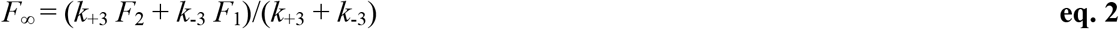

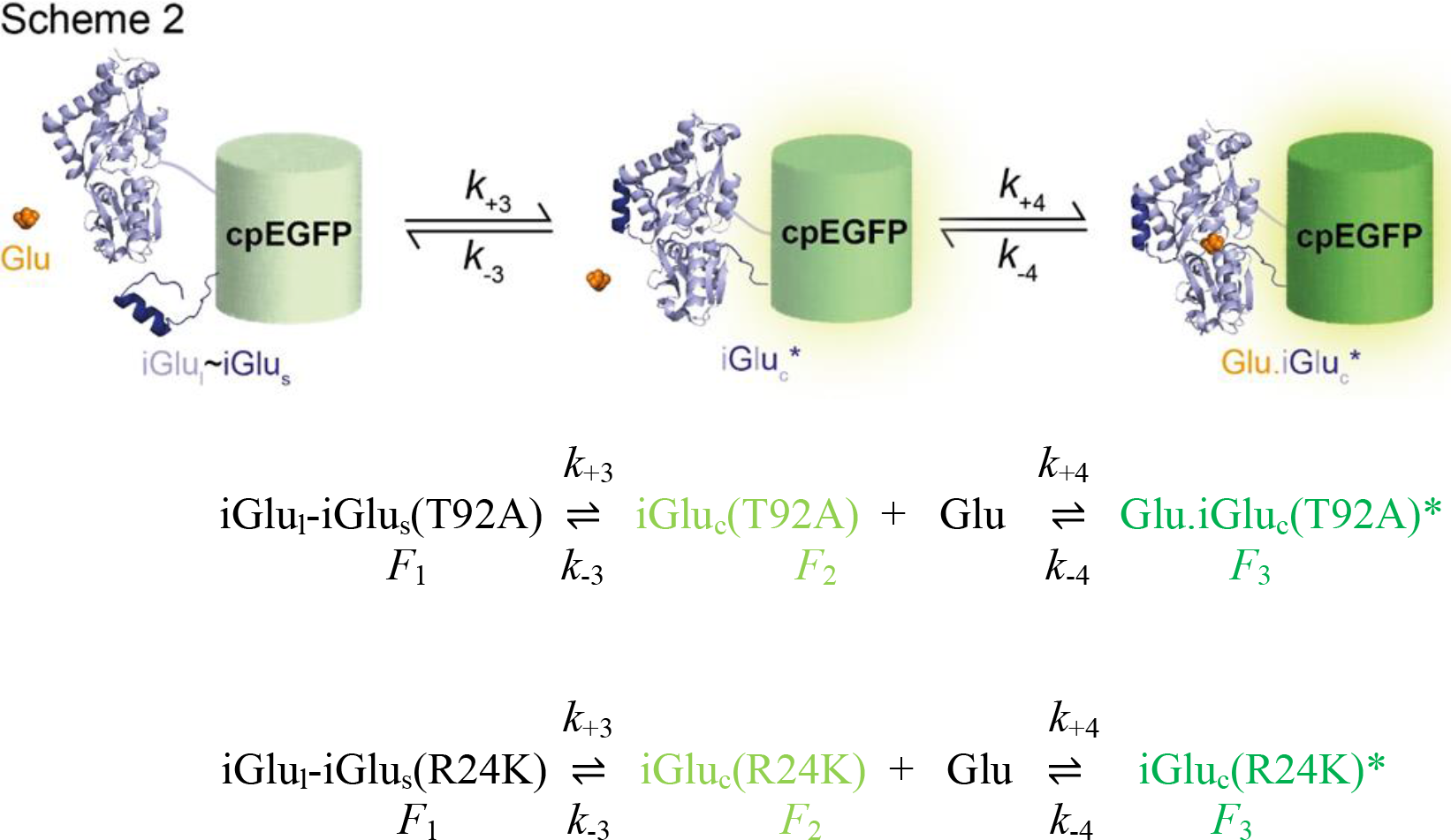

**Table 2.**
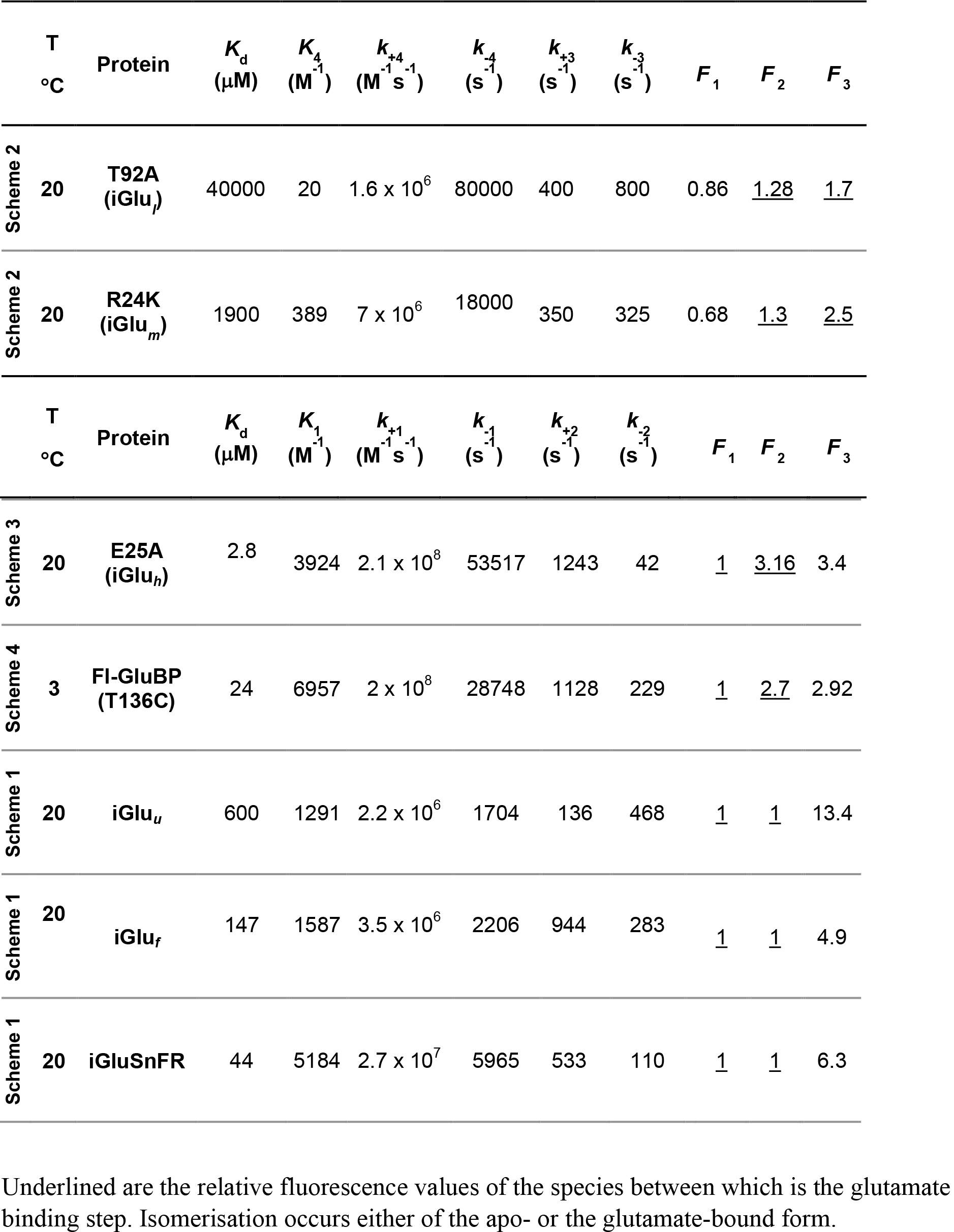
Kinetic parameters of iGlu_*l*_, iGlu_*m*_, iGlu_*h*_ and Fl-GluBP fitted to their respective mechanisms

It must be noted that the mechanisms denoted as **Scheme 1** and **Scheme 2** involve the same type of intermediates and represent two pathways leading to the same final product (see **Fig. 4G** in (3)).

### Kinetic mechanism of high affinity sensor iGlu-E25A (iGlu_h_)

The hyperbolic association rate plot of iGlu_*h*_ (**Fig. 1I,J**) appears similar to that of iGluSnFR and fast variant iGlu_*f*_ (3). However, two fluorescent states are apparent as an initial jump to higher fluorescence is followed by a measurable exponential process. To distinguish the two scenarios, the mechanism for iGlu_*h*_ is depicted in **Scheme 3**, in which rapid binding of glutamate induces a fluorescence enhancement which is then further increased by a subsequent isomerisation which stabilises the bound complex. An intermediate of ‘semi-complete’ (sc) structure iGlu_sc_(E25A) is thus postulated, which upon glutamate binding becomes more fluorescent Glu.iGlu_sc_(E25A). A further structural rearrangement then leads to the stable highly fluorescent complex Glu.iGlu_c_(E25A)* (**Scheme 3**). Dissociation occurred in a single exponential process with the measured rate of 15 s^−1^ (20 °C). Best fit parameters to the hyperbolic association rate plot described by **eq. 3** (20) were: *K*_1_ 3.92 × 10^3^ M^−1^, *k*_+2_ 1243 s^−1^ and *k*_−2_ 42 s^−1^, indicating strong stabilisation by the isomerisation, also explaining the single exponential dissociation kinetics (fitted *K*_d(overall)_ 2.8 μM, similar to measured 5.8 ± 0.2 μM). Global fitting gave *k*_+2_ of 2.1 × 10^8^ M^−1^s^−1^ indicating rapid, diffusion limited glutamate binding to Glu.iGlu_sc_(E25A). *F*_1_ has the value of 1, *F*_2_ is 3.16 and *F*_3_ is 3.4 (*F*_∞_ = 3.4).

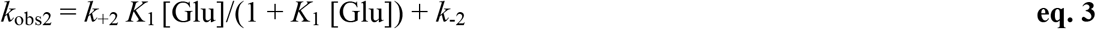

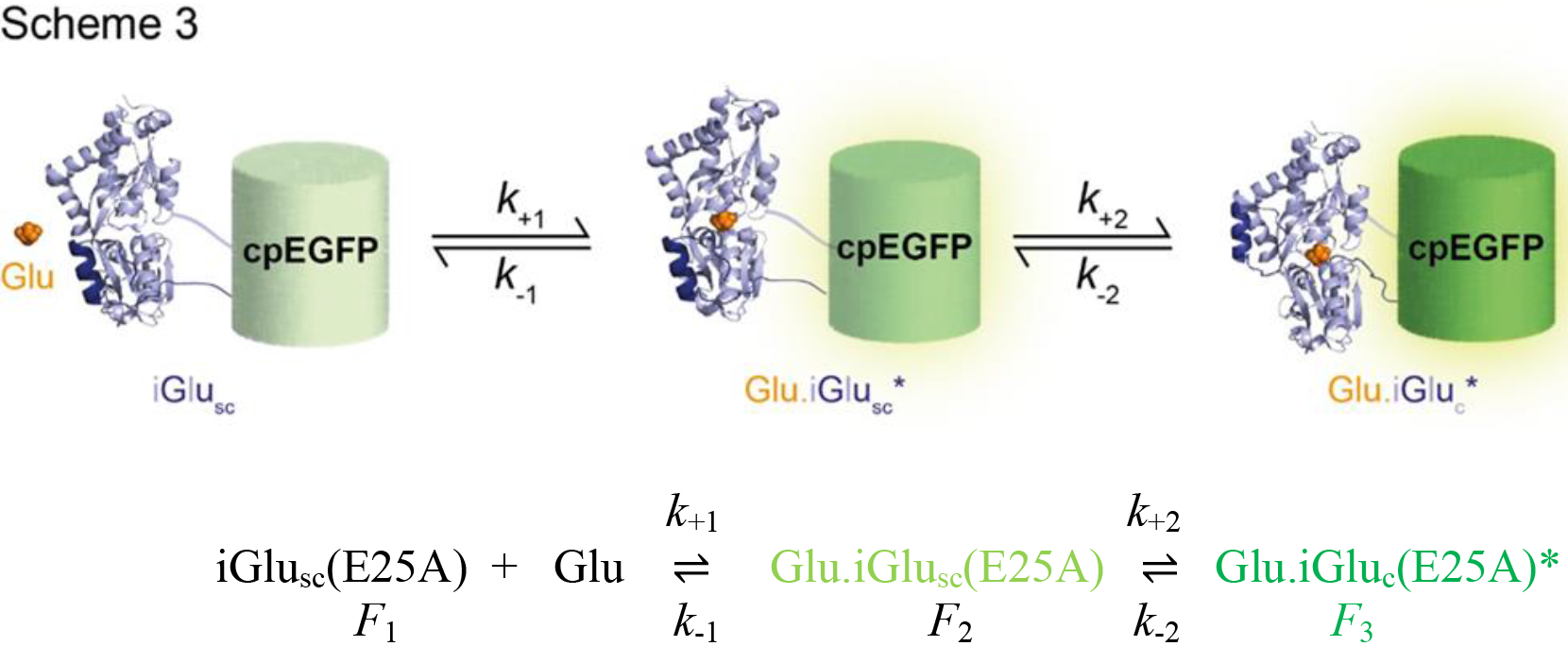

**Scheme 3** represents the same pathway as that in **Scheme 1** with the difference that in **Scheme 3**, the initial glutamate-bound intermediate has elevated fluorescence as well as the final product. Interestingly, phenomenologically a kinetically similar mechanism to that in **Scheme 3** was observed for a novel chemically labelled glutamate sensor, Fl-GluBP.

### Development and kinetic mechanism of Fl-GluBP

GluBP variants T71C, S72C, T83C, S90C, T92C, T136C, S137C, T140C and A210C were generated and covalently labelled with environmentally sensitive fluorophores acrylodan, BADAN, CPM, IDCC, IANBD ester and Oregon Green 488 maleimide. Fluorescently labelled derivatives were tested for fluorescence dynamic range, ligand selectivity and kinetic response to ligand binding. Of all the combinations tested, IANBD-T136C-GluBP (termed Fl-GluBP) stood out with a 2.9-fold fluorescence enhancement upon glutamate binding. The *K*_d_ of Fl-GluBP for glutamate was 9.7 ± 0.3 μM (**Fig. 1C** and **Table 1**), at physiological ionic strength, pH 7.5 and 20 °C, ~20- fold increased from the 600 nM reported for GluBP 600n (21).

The kinetics of the interaction of Fl-GluBP with glutamate were investigated by fluorescence stopped-flow at 3 °C (**Fig. 1L**). For association kinetic experiments, as the glutamate concentration increased, it became apparent that the measured single exponential rise is preceded by a jump to a level that represents most of the fluorescence increase, indicating that at saturating concentrations, the first phase of the interaction - ~ 90% of the fluorescence enhancement - was too fast to measure. Rates in the range of 200 to 800 s^−1^, showing saturation, were measured for the second phase, interpreted as an isomerisation. Plotting the isomerisation rate (*k*_obs_) as a function of glutamate concentration, a hyperbolic concentration dependence was observed (**Fig. 1M**). Dissociation kinetics were obtained by rapidly mixing saturated 0.5 μM Fl-GluBP (33 μM glutamate) with 457 μM GluBP 600n. A single exponential fluorescence decay at a rate of 217 ± 5 s^−1^ was observed at 3 °C (**Fig. 1N**), extrapolated to 1003 s^−1^ at 37 °C based on a linear Arrhenius plot (**Fig. 1N inset**).

The kinetic data measured at 3 °C for Fl-GluBP were interpreted in terms of a two-step mechanism in which rapid glutamate binding is followed by isomerisation (**Scheme 4**). Best fit parameters to the model were: for step 1, equilibrium constant *K*_1_ of 6957 M^−1^ and for the second step, *k’*_+2_ of 1228 s^−1^ and *k’*_−2_ of 229 s^−1^ were best fit giving *K*_d(overall)_ 24 μM (**Table 2**), comparable to the measured 10.6 ± 0.4 μM (**Fig. 2F** and **Table 3**). Global fitting of the data gave *k’*_+1_ of 2 × 10^8^ M^−1^s^−1^ and *k’*_−1_ of 28748 s^−1^, suggesting diffusion-limited glutamate binding. Taking the relative fluorescence (*F*_1_) value of Fl-GluBP as 1, Glu.Fl-GluBP* and Glu.Fl-GluBP^**^ have *F*_2_ of 2.7 and *F*_3_ of 2.92, respectively (*F*_∞_ = 2.9).

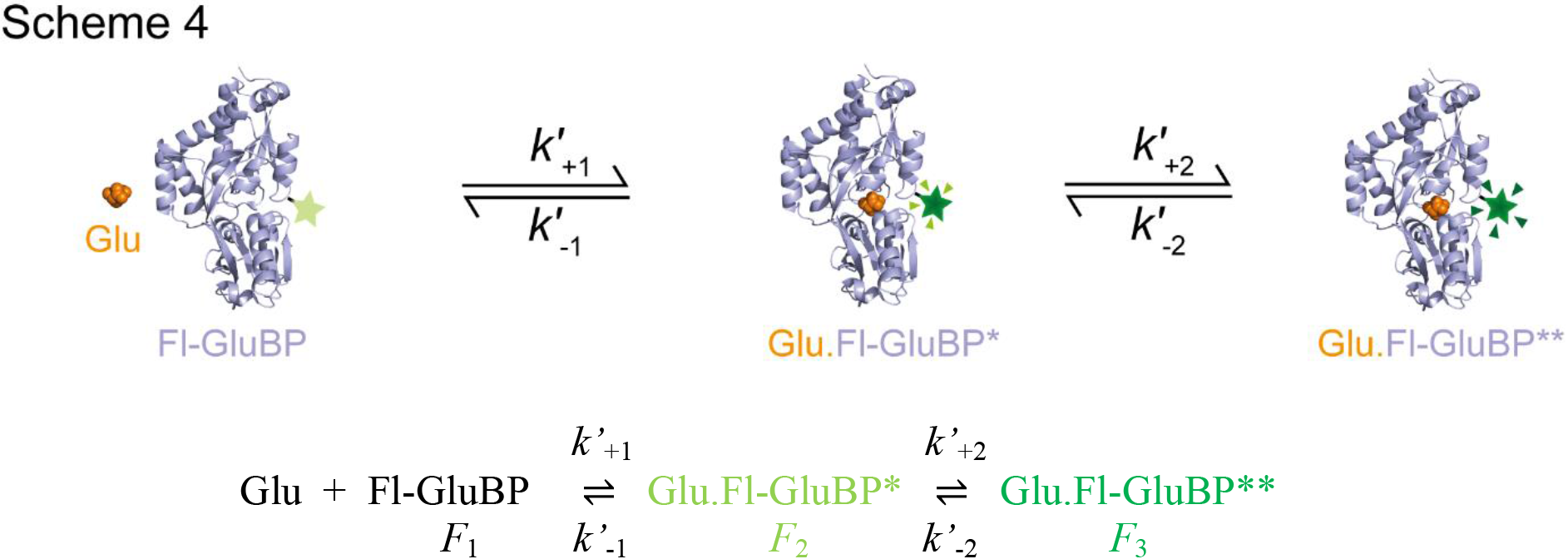

**Table 3.**
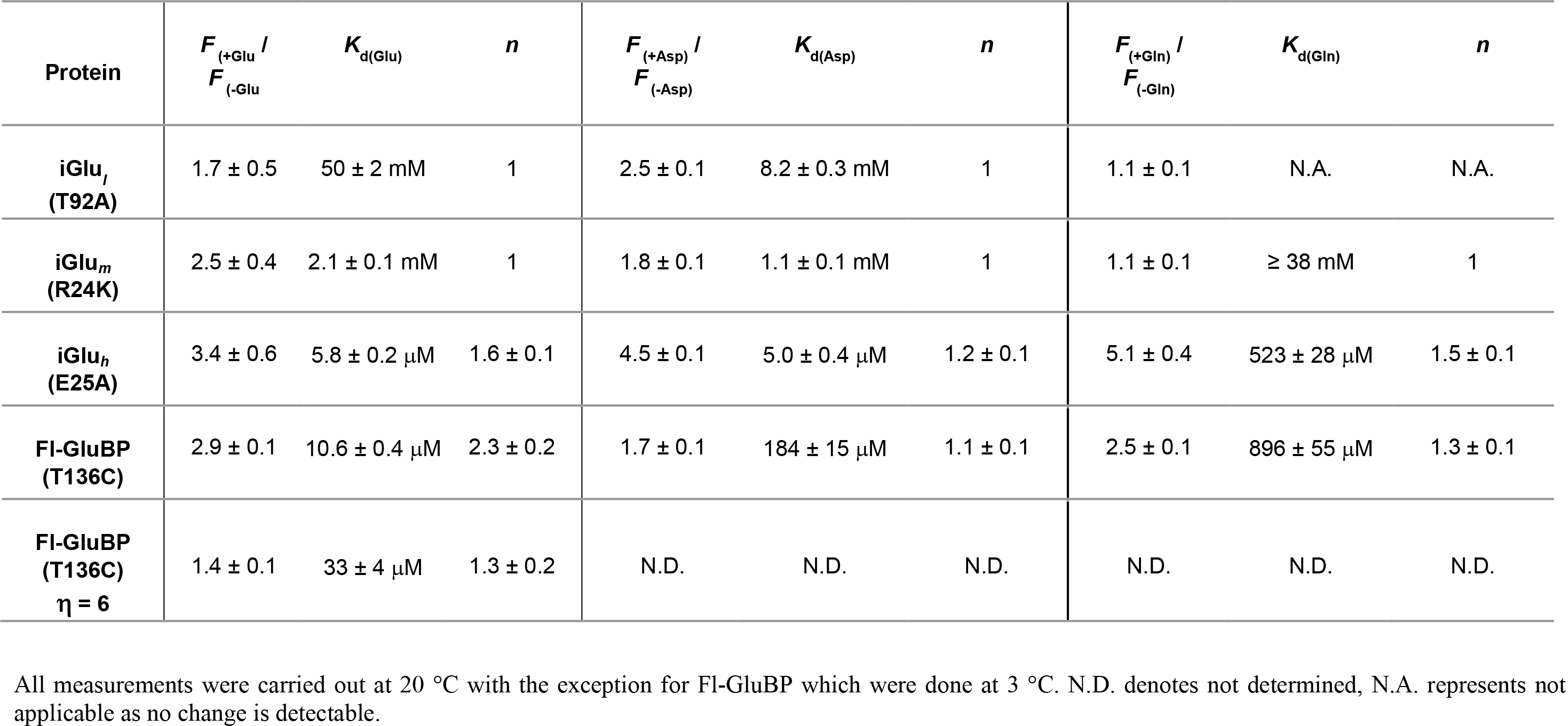
Affinity and selectivity of Fl-GluBP, iGlu_*h*_, iGlu_*m*_ and iGlu_*l*_ and for L-aspartate and L-glutamine.

**Figure 2.**
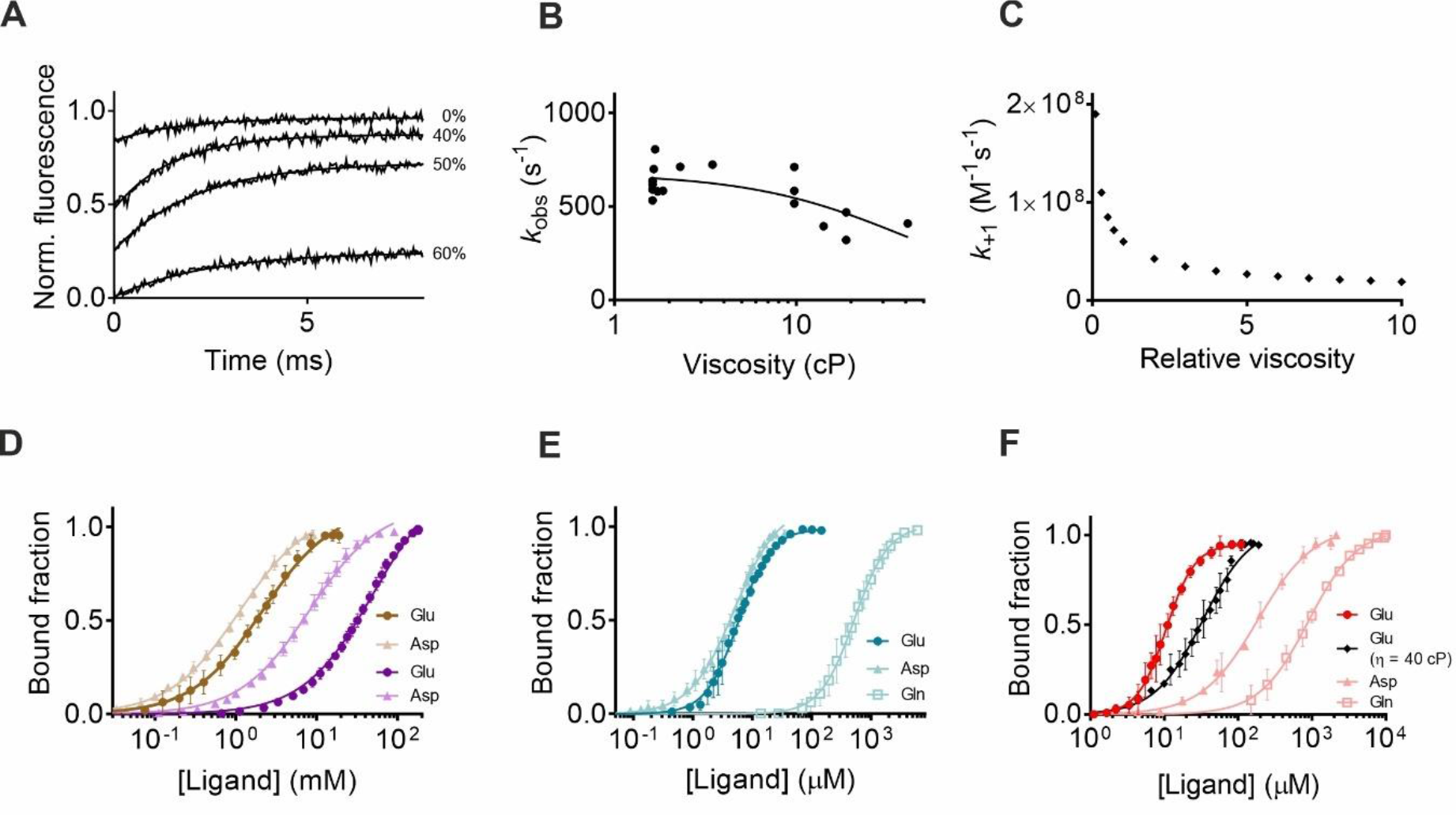
Viscosity dependence and selectivity. (**A**) Association kinetic records for Fl-GluBP in solvents with increasing viscosity at 3 °C. (**B**) Plot of observed association rates (*k*_obs_) against relative viscosity for Fl-GluBP. (**C**) Plot of predicted second order rate constant as a function of relative viscosity, at 3 °C. (**D**) Equilibrium titration of iGlu-T92A (iGlu_*l*_) (purple symbols and lines) and iGlu-R24K (iGlu_*m*_) (terracotta symbols and lines) with glutamate and aspartate at 20 °C. (**E**) Equilibrium titration of iGlu-E25A (iGlu_*h*_) with glutamate, aspartate and glutamine at 20 °C. (**F**) Equilibrium titration of Fl-GluBP with glutamate (in 0 and 60% glycerol) at 3 °C, aspartate and glutamine at 20 °C.

### Kinetics of glutamate binding to Fl-GluBP under increased viscosity

We measured the association kinetics of Fl-GluBP at increasing solvent viscosities to see if the diffusion-limited glutamate binding step is affected. Relative viscosity (η) was increased up to 6-fold. Interestingly, the amplitude of fluorescence enhancement rather than the rate of binding was affected. Holding [glutamate] at 50 μM (in the mixing chamber), the rapid binding step appeared as a progressively smaller ‘jump’ in fluorescence intensity, and was completely abolished in 60% glycerol (**Fig. 2A**). The disappearance of the initial fast fluorescence was the result of the apo-state fluorescence intensity increasing with greater viscosity. Increasing solvent viscosity thus had a similar effect on the fluorescence intensity of apo-Fl-GluBP to glutamate binding. At relative viscosity η of 6, the glutamate binding invoked fluorescence increase purely reflected the conformational change of the protein.

In contrast, the rate of isomerisation was slowed down to 400 s^−1^ at 50-60% glycerol from 600 s^−1^ (**Fig. 2B**). Applying a previously developed theory (22) from the foundation laid down in (23) for the effect of solvent viscosity on first order processes, we obtained a good fit for our isomerisation data to **eq. 4**, where C and σ (σ has units of viscosity) are adjustable parameters, η is solvent viscosity. The pattern of the rate plot fits well to the theory that, in the viscosity range examined, both internal friction of the protein and friction of the molecule with the solvent contribute to decreasing the rate constant (22) (**Fig. 2B**). The fitted values gave C = 1.96 × 10^8^ cP s^−1^, σ = 40.45 cP and *E*_o_ = 4.87 kcal mol^−1^. The small activation energy indicates that most of the change in the rate constant is due to the change in viscosity (22).

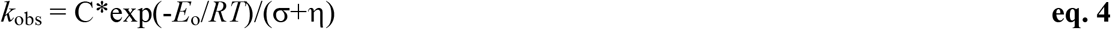

### Ligand selectivity of iGlu_*l*_, iGlu_*m*_, iGlu_*h*_ and Fl-GluBP

Each of the mutations giving iGlu_*l*_, iGlu_*m*_, iGlu_*h*_ shifted the selectivity towards aspartate which does not disqualify these variants from investigations at excitatory synapses in the hippocampus where glutamate fully accounts for neurotransmission (24). The selectivity for glutamate over glutamine is a more complex issue. The lowest affinity variants iGlu_*l*_ and iGlu_*m*_ appear to show the highest selectivity for glutamate, their response to glutamine is hardly detectable. Fl-GluBP, similarly to iGlu_*h*_ is highly selective for glutamate (*K*_d_ 10.6 ± 2.3 μM) over aspartate (*K*_d_ 184 ± 15 μM) and glutamine (*K*_d_ 896 ± 55 μM), with smaller, 1.7 and 2.5-fold fluorescence enhancements, respectively (**Fig. 2F** and **Table 3**). However, in an environment where both ligands are present, iGlu_*h*_ and Fl-GluBP could yield composite signals due to their affinities being relevant to the physiological concentration ranges and high fluorescence dynamic range for both ligands (**Fig 2D,E** and **Table 3**). At 100 μM glutamate for example, Fl-GluBP would be saturated by while giving only 10% of the signal at 100 μM glutamine. However, Fl-GluBP would function well in the synaptic cleft where glutamine concentration is negligible. Fl-GluBP may also be suitable for detecting glutamine in an environment where glutamate concentration is negligible. D-Ser, glycine and GABA do not evoke any fluorescence response from Fl-GluBP (data not shown).

## DISCUSSION

Protein-based fluorescent glutamate sensors have the potential for real-time monitoring of synaptic and cellular glutamate concentration changes. We have developed both genetically encoded and chemically labelled fluorescent glutamate sensors and characterised the kinetic mechanisms of their glutamate sensing. Previously described iGlu_*f*_ and iGlu_*u*_ responded to glutamate binding with a single exponential fluorescence increase which occurred in an isomerisation step following glutamate binding (**Scheme 1**) (3). Novel variants, iGlu_*l*_ and iGlu_*m*_ with mM affinity for glutamate, follow an alternative kinetic path whereby reattachment of GluBP fragments, accounting for part of the fluorescence enhancement, is required for glutamate to bind to the reformed complex, which causes further fluorescence enhancement (**Scheme 2**). For iGlu_*l*_ and iGlu_*m*_ the conformer with the separate large and small GluBP fragments is in an equilibrium with the reassembled, ‘complete’ GluBP (a low-fluorescence conformer) to which glutamate preferentially binds followed by most of the fluorescence enhancement. It must be noted that with the exception of iGlu_*l*_ and iGlu_*m*_, equilibrium titration curves are best fit to the Hill equation giving Hill coefficient (*n*) values of 1.6 to 2.3. However, as the kinetic analyses reveal, none of the kinetic data indicate cooperativity of binding. Therefore, *n* =1 is used in the kinetic analyses.

For iGlu_*h*_, glutamate binding is followed by isomerisation (**Scheme 3**). Fluorescence enhancement of iGlu_*h*_ is biphasic with most of the increase occurring in the diffusion limited glutamate binding phase, indicating that apo-GluBP in this variant exists in a ‘semi-complete’ conformation in which the small fragment may already be attached to the large fragment albeit not in a stable conformation.

Fl-GluBP, a glutamate sensor generated by fluorescent derivatisation of GluBP with a synthetic fluorophore also yields most of its fluorescence enhancement in the initial glutamate binding phase which occurs with a diffusion limited rate constant (**Scheme 4**). A subsequent isomerisation further stabilises the fluorescent complex. The rate of isomerisation for Fl-GluBP is fitted to saturate at 2220 s^−1^ at 3 °C, indicating that Fl-GluBP will be suitable as a real-time tracker of synaptic glutamate transients.

While we were therefore unable to measure the second order rate constant (*k*_+1_) for glutamate binding even at increased solvent viscosity, we attempt to estimate it. An inverse relationship between rate constant and viscosity has been reported (25). Reasoning, that for it to be too fast to measure at 3 °C and have a relative viscosity of 6 at 50 μM glutamate, the observed association rate needs to be > 1000 s^−1^, *k*_+1_ > 2 × 10^7^ M^−1^s^−1^ is required. At relative viscosity of 1, *k*_+1_ is predicted to be 10-fold higher, > 2 × 10^8^ M^−1^s^−1^ at 3 °C (**Fig. 2C**). Assuming a 2-fold increase for every 5 °C increase in temperature, *k*_+1_ > 2 × 10^9^ M^−1^s^−1^ at 20 °C and *k*_+1_ > 3 × 10^10^ M^−1^s^−1^ 37 °C are predicted. If another empirical formula in which there is an inverse relationship between diffusion-controlled reaction rate constant and the square root of relative viscosity is used (26), the estimates are in the same range. These values are consistent with diffusion limited glutamate association kinetics which makes Fl-GluBP and iGlu_*h*_ potential real-time detectors of synaptic glutamate release kinetics.

There is strong structural homology between GluBP and the S1S2 glutamate binding domain of AMPAR. Neither iGluSnFR-type probes, nor the previously studied Trp fluorescence changes allowed the observation of glutamate binding itself. The observed conformational changes by Trp fluorescence led to the Venus fly-trap model for ligand binding to bacterial periplasmic binding proteins (14,27) and the homologous S1S2 glutamate binding construct derived from AMPAR (15) which does not explain the rapid opening of the AMPAR ion channel triggered by ligand glutamate binding. Neither can the iGlu_*u*_ fluorescence response, which is based on a protein conformational change, as explained above. The kinetic examination of Fl-GluBP and iGlu_*h*_, however, reveals that they signal diffusion limited binding of glutamate. We propose that a similar mechanism of diffusion-limited glutamate binding exists for and forms the basis of rapid gating of AMPAR.

Moreover, a fluorescent sensor like Fl-GluBP may be useful for measuring synaptic viscosity. This may be through the relative fluorescence measurements of the binding and isomerisation steps or by lifetime imaging as the lifetime is expected to increase if a singlet-exciplex intermediate is formed (28,29).

The two low-affinity glutamate sensors iGlu_*l*_ and iGlu_*m*_ have fast *off*-rates of 800 s^−1^ (fitted value) and 365 s^−1^ (measured at 20 °C), respectively. The *off*-rate for Fl-GluBP is measured as 217 s^−1^ at 3 °C, the extrapolated value at 37 °C is 2000 s^−1^, from its Arrhenius plot. All three sensors thus allow monitoring processes on the sub-millisecond time scale, at the temperatures of physiological experiments (34 - 37 °C). Through their broad affinity range and mechanistic variety, the above genetically encoded and chemically labelled fluorescent glutamate sensors could form part of a toolkit designed for monitoring the different processes that glutamate undergoes in neurotransmission and cellular homeostasis.

## CONCLUSION

Subtle structural changes brought about by single amino acid substitutions, in addition to affecting their ligand binding affinity and selectivity, also redirect the kinetic paths of the fluorescence response of iGluSnFR variants. A novel probe labelled with synthetic fluorophore reveals diffusion limited glutamate binding, hints at the AMPAR response mechanism and may be suitable for measuring synaptic viscosity.

## Author Contributions

C.C. and S.K generated the genetically encoded proteins, performed experiments, analysed data and generated figures; N.H. generated the chemically labelled probes. K.T. designed the project and wrote the paper.

## Acknowledgements

This work was funded by Wellcome Trust grant 094385/Z/10/Z and BBSRC grants BB/M02556X/1 and BB/S003894/1 to K.T. Dr Alamin Mohammed is thanked for comments on the manuscript. Melissa Matthews is thanked for her help with the characterisation of the R24K iGluSnFR variant.

